# A comparison of tools for copy-number variation detection in germline whole exome and whole genome sequencing data

**DOI:** 10.1101/2021.04.30.442110

**Authors:** Migle Gabrielaite, Mathias Husted Torp, Malthe Sebro Rasmussen, Sergio Andreu-Sánchez, Filipe Garrett Vieira, Christina Bligaard Pedersen, Savvas Kinalis, Majbritt Busk Madsen, Christina Westmose Yde, Lars Rønn Olsen, Rasmus L. Marvig, Olga Østrup, Maria Rossing, Finn Cilius Nielsen, Ole Winther, Frederik Otzen Bagger

## Abstract

**Background:** Copy-number variations (CNVs) have important clinical implications for several diseases and cancers. The clinically relevant CNVs are hard to detect because CNVs are common structural variations that define large parts of the normal human genome. CNV calling from short-read sequencing data has the potential to leverage available cohort studies and allow full genomic profiling in the clinic without the need for additional data modalities. Questions regarding performance of CNV calling tools for clinical use and suitable sequencing protocols remain poorly addressed, mainly because of the lack of good reference data sets.

**Methods:** We reviewed 50 popular CNV calling tools and included 11 tools for benchmarking in a unique reference cohort encompassing 39 whole genome sequencing (WGS) samples paired with analysis by the current clinical standard —SNP-array based CNV calling. Additionally, for nine of these samples we performed whole exome sequencing (WES) performed, in order to address the effect of sequencing protocol on CNV calling. Furthermore, we included Gold Standard reference sample NA12878, and tested 12 samples with CNVs confirmed by multiplex ligation-dependent probe amplification (MLPA).

**Results:** Tool performance varied greatly in the number of called CNVs and bias for CNV lengths. Some tools had near-perfect recall of CNVs from arrays for some samples, but poor precision. Filtering output by CNV ranks from tools did not salvage precision. Several tools had better performance patterns for NA12878, and we hypothesize that this is the result of overfitting during the tool development.

**Conclusions:** We suggest combining tools with the best recall: GATK gCNV, Lumpy, DELLY, and cn.MOPS. These tools also capture different CNVs. Further improvements in precision requires additional development of tools, reference data sets, and annotation of CNVs, potentially assisted by the use of background panels for filtering of frequently called variants.

## Background

Large cohort-based genome-wide association studies provided us with the tools and knowledge to understand numerous phenotypic traits and diseases by single nucleotide polymorphisms (SNPs) and short (<50 bp) insertions and deletions in the genome. It is, however, more challenging to assess the role of the larger structural variations which have proven to b e important for the regulation and function of many gene products. This is particularly the case for copy number variations (CNVs) [1] which were first described in healthy humans [2] but have since been associated with diseases, especially neurodevelopmental disorders and cancer [3, 4]. CNVs are estimated to contribute to 4.8 –9.5% of the genome and one or multiple exon copy number changes can affect gene expression levels or induce chromosomal rearrangements causing various disorders and diseases [5, 6].

The current clinical standard method for CNV assessment remains array-based CNV identification, either from array-based comparative hybridization or SNP-array approaches [7, 8]. While these arrays provide relatively accurate, cost-effective, and precise identification of CNVs, the use of short-read sequencing (or next generation sequencing, NGS), is not limited to the specific regions included on the arrays, has higher potential to identify novel CNVs, and has higher resolution at predicting both the breakpoints and shorter CNVs [9]. Long-read sequencing is still cost-preventive for routine diagnostics and uniquely suited for structural variants (SVs). The alternative—NGS—bears the potential of a single assay for complete genomic analysis that allows for automated identification of both SNPs and structural variants from the same data. CNV calling from NGS does, however, create new challenges; such as dealing with variable coverage across the genome, alignment bias for deletions, and read-length limit and insensitiveness towards repetitive and breakpoint regions [10]. Furthermore, short-read sequencing increases mapping ambiguity consequently increasing the complexity of CNV detection [10]. This is particularly true for whole exome sequencing (WES) or targeted gene panels, as the sequencing coverage and read depth in different areas are highly variable.

CNV calling algorithms can be based on one or more approaches: read-pair (RP), read-depth (RD), split read (SR), or assembly (AS) algorithms [11] (Figure 1). Most CNV calling tools are based on RD algorithms predicting CNVs from the changes based on read coverage in different areas of the genome.

**Figure 1.**
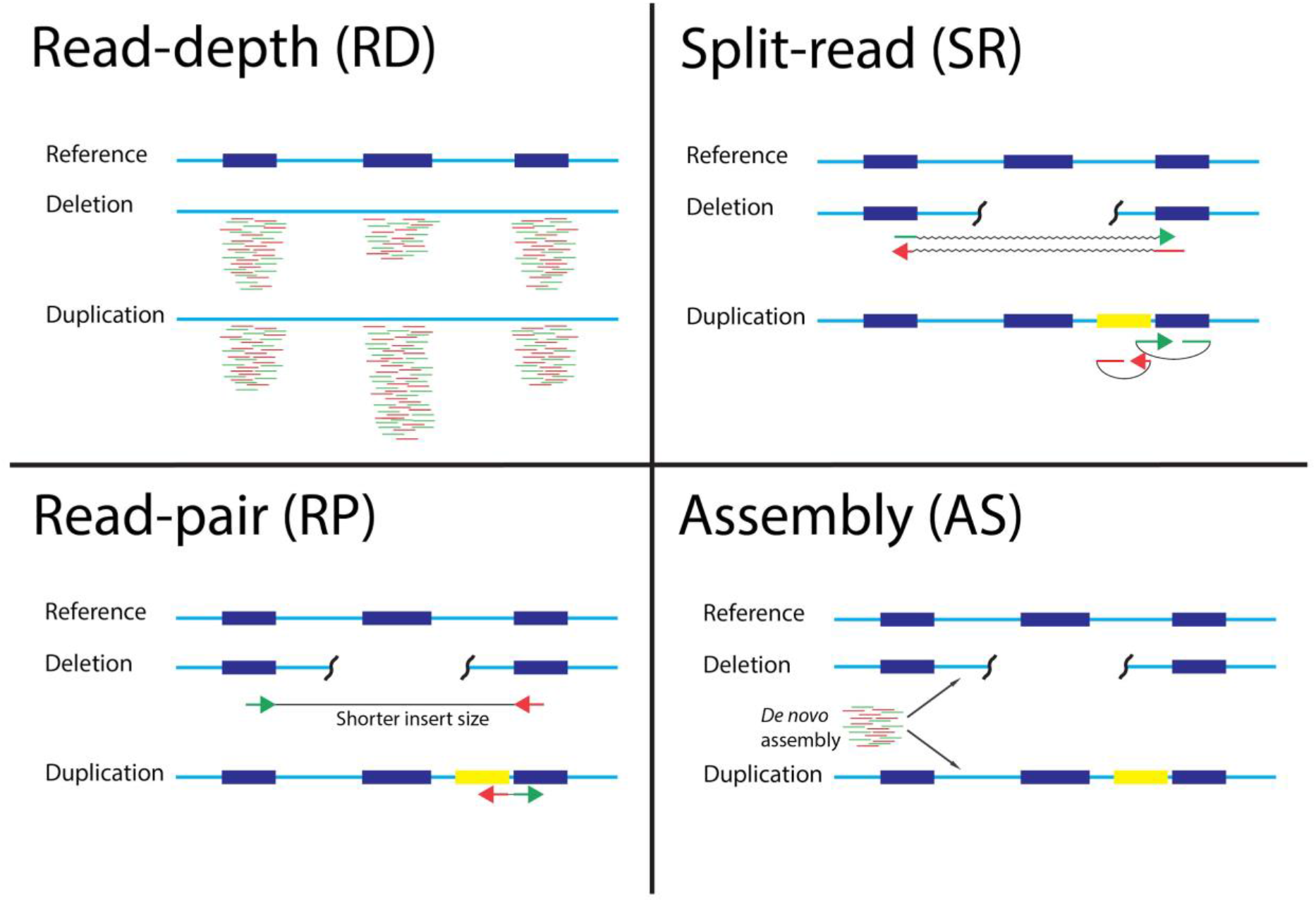
Schematic visualization of different approaches for calling CNVs from NGS data. RD detects local difference in read-depth, SR detects unmatched read pairs, RP detects decreased insert size or swapped read directions between read pairs, and AS performs *de novo* assembly to best explain read distribution.

With the increasing attention for the identification of SVs, including CNVs, and their clinical use, multiple SV databases have been established, such as CNV tracks in the UCSC database [12, 13] or gnomAD SV [14]. These databases help to evaluate predicted CNVs more accurately; however, the correct CNV identification itself remains challenging. In recent years, a number of tools for calling CNVs from NGS sequencing data, have been developed [11]. Currently, there is no clear standard tools for SV detection (including CNV) and a lack of a comprehensive benchmark of the tools on known NGS datasets using more than a single sample [9, 11, 15] and no clear standard tools for SV (including CNV) detection.

Here, we report an evaluation of 11 CNV detection tools for NGS and aim to identify the most reliable and clinically applicable software, whether based on WES or whole genome sequencing (WGS). We used the standard CytoScan HD SNP-array as a reference method for CNV detection and assessed the potential increase in sensitivity of WGS in comparison to WES. We Selected the best performing tools and attempted to optimize the CNV calling to improve precision. Finally, We Suggest a combination of four tools (GATK gCNV, Lumpy, DELLY, and cn.MOPS) for a balanced CNV recall and precision.

## Results

### Review of 50 most popular CNV calling tools

We reviewed 50 most popular tools for CNV calling (Figure 2, Supplementary Table 1). The tools were included in the benchmark if they were: (1) developed for calling CNVs from WES or WGS data, (2) developed for germline CNV calling, (3) recently developed or highly cited (>100 citations as of March, 2019, using the number of citations as the only available proxy for popularity of use), (4) still maintained, and (5) the latest versions of the tool was more recent than 5 years. However, several tools passing all these criteria were not suitable for inclusion in this benchmark (detailed information in Supplementary Table 1). After applying the selection criteria, 11 tools (in bold in Figure 2) were selected for further performance evaluation.

**Figure 2.**
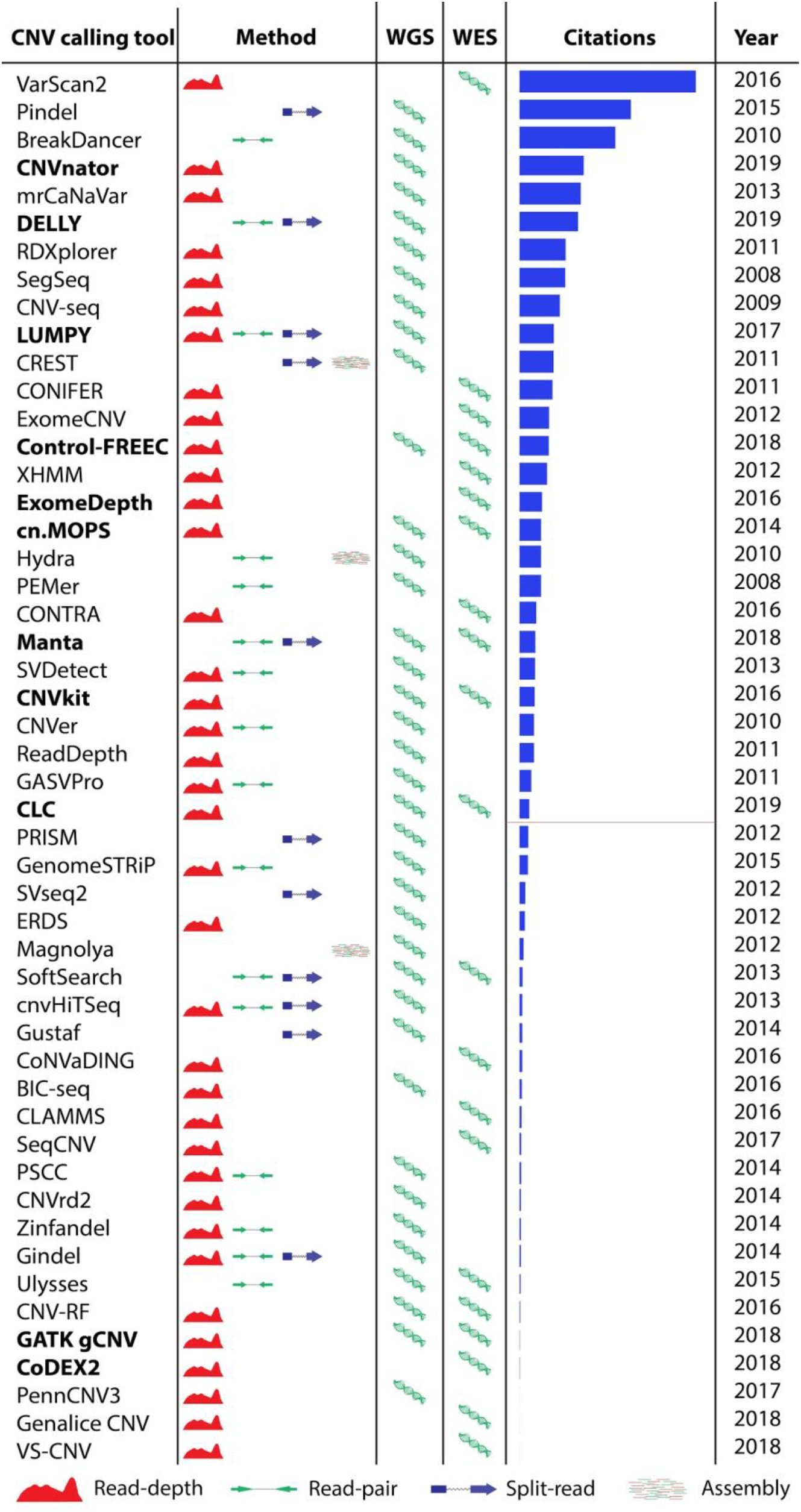
Overview of methods CNV calling tools applies, input NGS data, citation number from Google Scholar and available latest version for each tool as of March, 2019. Tools highlighted with bold font are included in the benchmark, the horizontal red line shows the cutoff for the citation number.

### Datasets used for the benchmark study

The datasets used for the benchmark are listed in Table 1. Gold Standard sample NA12878 from 1000 Genomes Project was used with the CNVs which were published by Haraksingh *et al*., 2017 [8]. For the in-house GB01–GB08 and GB09–GB38 samples the true reference was considered to be Nexus software-produced filtered CNV calls from CytoScan HD SNP-array, which have previously been shown to be among the best-performing array platforms [8, 16]. To account for imperfections in the SNP-array CNV calling, we compared all CNV calls made by different CNV calling tools (Figure 5A and Figure 5B). Furthermore, selected CNVs were confirmed for the GB40–GB51 samples using multiplex ligation-dependent probe amplification (MLPA).

**Table 1.**
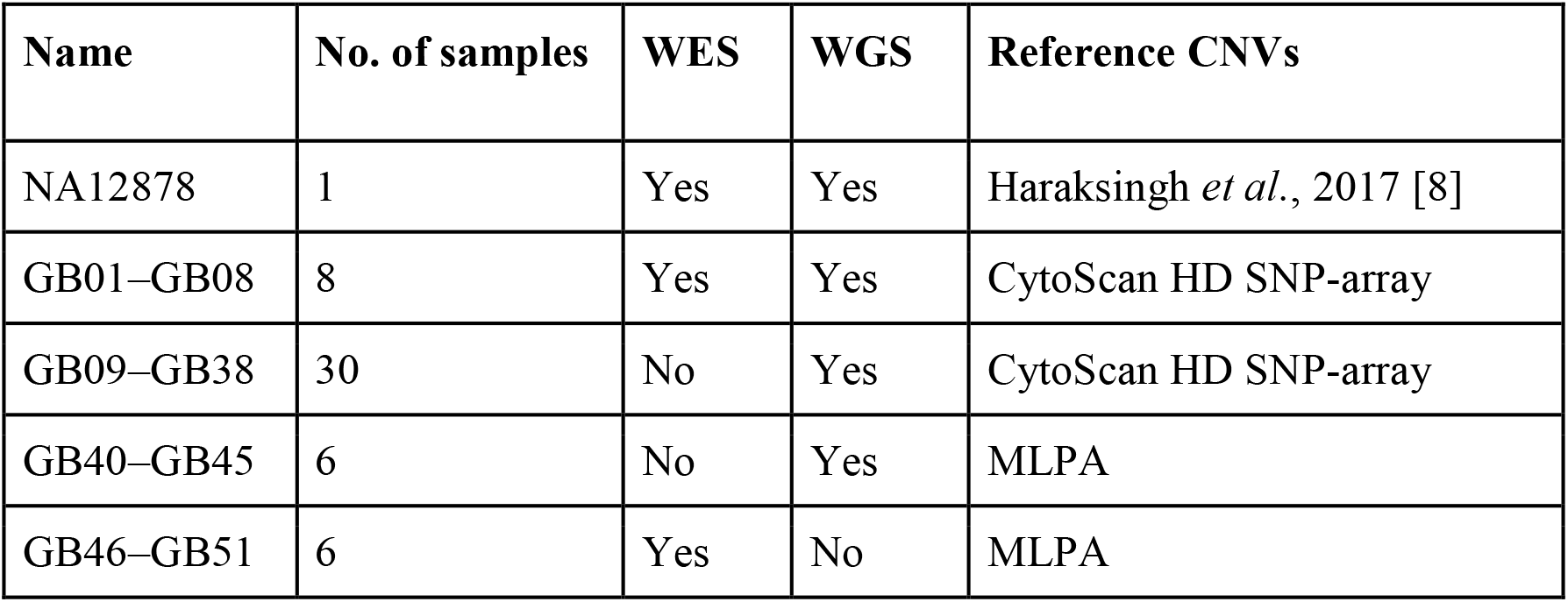
Datasets used in this benchmark study.

| Name | No. of samples | WES | WGS | Reference CNVs |
| --- | --- | --- | --- | --- |
| NA12878 | 1 | Yes | Yes | Haraksingh *et al.*, 2017 [8] |
| GB01–GB08 | 8 | Yes | Yes | CytoScan HD SNP-array |
| GB09–GB38 | 30 | No | Yes | CytoScan HD SNP-array |
| GB40–GB45 | 6 | No | Yes | MLPA |
| GB46–GB51 | 6 | Yes | No | MLPA |

**Figure 5.**
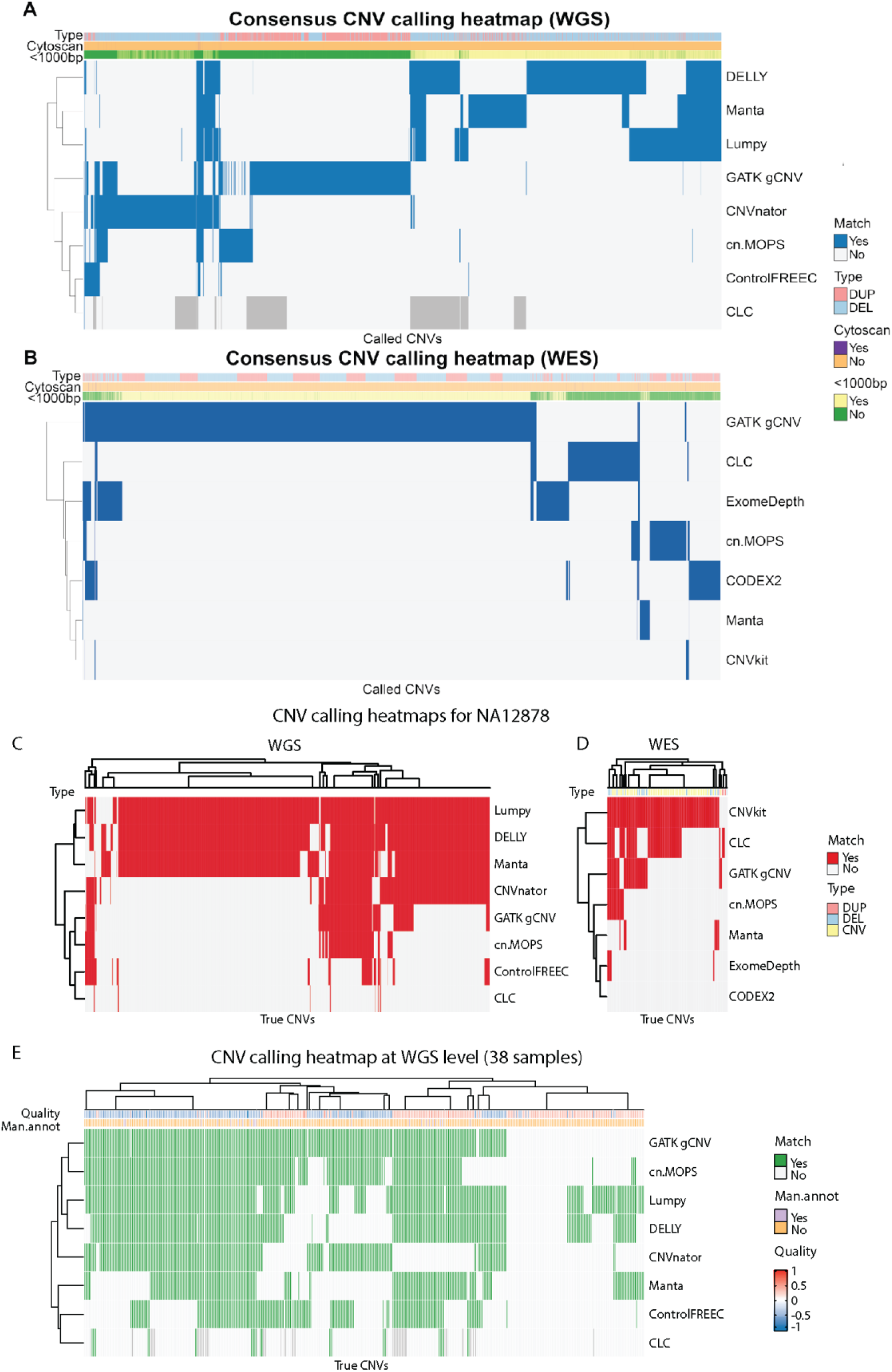
Heatmap showing all called CNVs across all samples (A–B) and called CNVs overlap with the true CNVs (C–E). (A) WGS (*n* = 407,671) and (B) WES level (*n* = 9,944). Each row represents a tool and a blue field denotes a call of the given CNV. All CNVs from each sample were merged across tools, such that any overlapping calls of either duplications or deletions were combined to one. Blue color denotes that the given CNV was called by the tool. The order of rows/columns for WES data and rows for WGS data was determined using complete-linkage hierarchical clustering with Euclidean distance, while the order of columns for WGS data was determined using a combination of k-means and hierarchical clustering due to memory restrictions. Darker grey coloring (WGS only) indicates that the tool was not run for the sample which contained the CNV. (C) 2,076 WGS-based and (D) 81 WES-based true CNVs in NA12878 sample. The order of rows/columns was determined using complete-linkage hierarchical clustering with Euclidean distance. (E) CNV calling heatmap for 471 true CNVs at and WGS level in 38 samples (GB01–38). Column dendrogram shows clustering to the level of 20 clusters to reduce complexity. The Quality annotation represents the probe median score from CytoScan HD SNP-array and the Man.annot. refers to whether the CNV was independently manually confirmed. A positive quality score corresponds to duplications, and negative scores denote deletions. Darker grey coloring indicates that the tool was not run for the sample which contained the CNV. The order of rows/columns was determined using complete-linkage hierarchical clustering with Euclidean distance.

### 11 CNV calling tools included in the benchmark

We selected 11 tools for the benchmark. Eight tools use a read depth approach: CNVnator [17], CLC Genomics Workbench [18], GATK gCNV [19], cn.MOPS [20], ExomeDepth [21], CNVkit [22], CoDEX2 [23] and Control-FREEC [24]. Other SV callers that include CNVs, such as DELLY [25], Manta [26] and LUMPY [27] use a combined approach and apply more than one CNV calling algorithm for more accurate predictions (Figure 2; detailed tool algorithms are provided in Supplementary File 1). All tools were run using the default parameters and following author recommendations when available. Most CNV calling tools are developed for either WES or WGS. However, cn.MOPS, CLC Genomics Workbench, CNVkit, Manta, and GATK gCNV tools are capable of calling germline CNVs using both WES and WGS data. For the NA12878 Gold Standard sample we also report CNVs called by Haraksingh *et al*., 2017 [8] using the consensus of several methods, including non-NGS based approaches.

### CNV length and type distribution for CNV calling tools

All tools called more deletions than duplications for NA12878 Gold Standard sample (Figure 3A). However, the total number of called CNVs varied greatly between the tools. GATK gCNV called more duplications and deletions in both WES and WGS samples compared to other tools. CODEX2 called the lowest number of deletions and duplications in WES samples and CLC Genomics Workbench called the lowest number of CNVs in WGS data. None of the CNV calls were filtered with a cutoff on confidence metrics, except where it was recommended by the authors of the tool (CODEX2) or the filtered files were created automatically (CLC Genomics Workbench). CODEX2 called a similar number of CNVs as provided in NA12878 Gold Standard true CNV set, but this did not equate that all the CNV calls were true positives. Similar patterns were observed for GB01–GB38 samples (Supplementary Figure 1).

**Figure 3.**
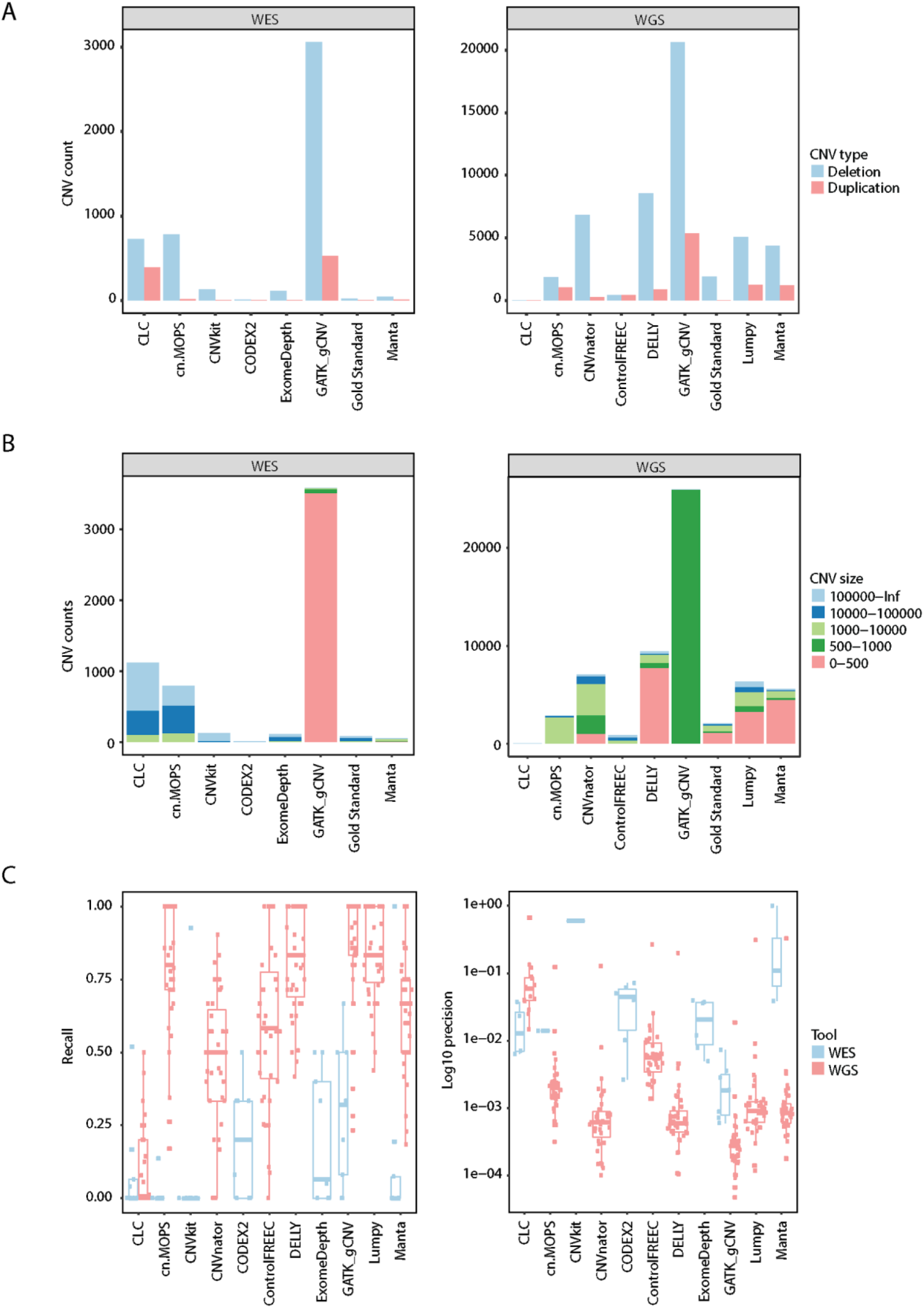
(A) Number of duplications and deletions called by CNV calling tools in WES and WGS data for the NA12878 sample. (B) Number CNVs called by all tools in WES and WGS data for the NA12878 sample colored by length. (C) Box plots and scatter plots for recall and precision results for 11 CNV calling tools.

CNV calls differed in lengths and frequencies among the tools in WES and WGS of NA12878 sample (Figure 3B). CLC Genomics Workbench and cn.MOPS called a high number of CNVs longer than 10,000 bp while GATK gCNV called mainly CNVs shorter than 500 bp in WES and 500–1,000 bp in WGS. GATK gCNV called shorter CNVs than any other tool. Furthermore, cn.MOPS, CNVnator, and Control-FREEC predicted more >1,000 bp length CNVs than other tools for WGS NA12878 sample. Half of CNVs in NA12878 were shorter than 500 bp as per Gold Standard truth CNV set. Similar patterns were observed for GB01 – GB08 WES and GB01–GB38 WGS samples (Supplementary Figure 2).

### Precision and recall of CNV calling tools

Given CNVs from CytoScan HD SNP-array for GB01–GB38 samples and NA12878 Gold Standard truth CNV set, true positive (TP), true negative (TN), false positive (FP), and false negative (FN) CNV calls were identified for each of 3 9 samples for WGS, and 9 samples (GB01–GB08 and NA12878) for WES (Supplementary Table 2). The criteria of an overlap of 1 bp between the Cytoscan HD SNP-array called CNV and the NGS-based tool CNV call was used for the CNV call to be classified as a TP. Re call and precision were calculated using the following formulas: 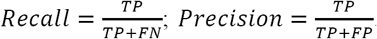. GB01–GB38 samples had a total 2–103 (median 7) CNVs called by CytoScan HD SNP-array, whereas 2,076 CNVs were called for NA12878 from the Gold Standard truth CNV set. For WES data, only CNVs covering exons were considered (1–63 (median 4) for GB01–GB08 and 233 for NA12878).

GATK gCNV recall was best for both WES and WGS data (Figure 3C), followed by Lumpy, DELLY, cn.MOPS, and Manta. All tools performed poorly on the WES dataset. While recall for WGS in all tools, except CLC Genomics Benchmark, was fair, precision was lacking for all the tools, with a maximum precision of 66.7% (Figure 5B). Tools that called a higher total number of CNVs, also had higher recall, but lower precision. The only two tools which use CNV call filtering (CLC Genomics Workbench and CODEX2) had a low recall compared to the tools which did not filter CNVs as part of their default settings. Collectively, recall approached 1 for several tools, but came at the expense of precision, which was lower than 31% in WGS data for the four best recalling tools (GATK gCNV, Lumpy, DELLY, and cn.MOPS).

### CNV call filtering possibilities for CNV calling tools

To explore the possibility of filtering CNV calls to improve precision we analyzed recall and precision at sliding confidence cutoff values. Briefly, for each tool, we calculated recall and precision at different thresholds defined by the percentiles of the tool-defined confidence metric (see Methods section for details).

The recall on WES was low, regardless of filtering, and that precision generally did not improve as a function of the threshold, and could not be easily interpreted as being asymptotically negative (Figure 4). The exception to this was CNVkit, which displayed high recall and precision on the WES NA12878 sample and offered the possibility of meaningful filtering.

**Figure 4.**
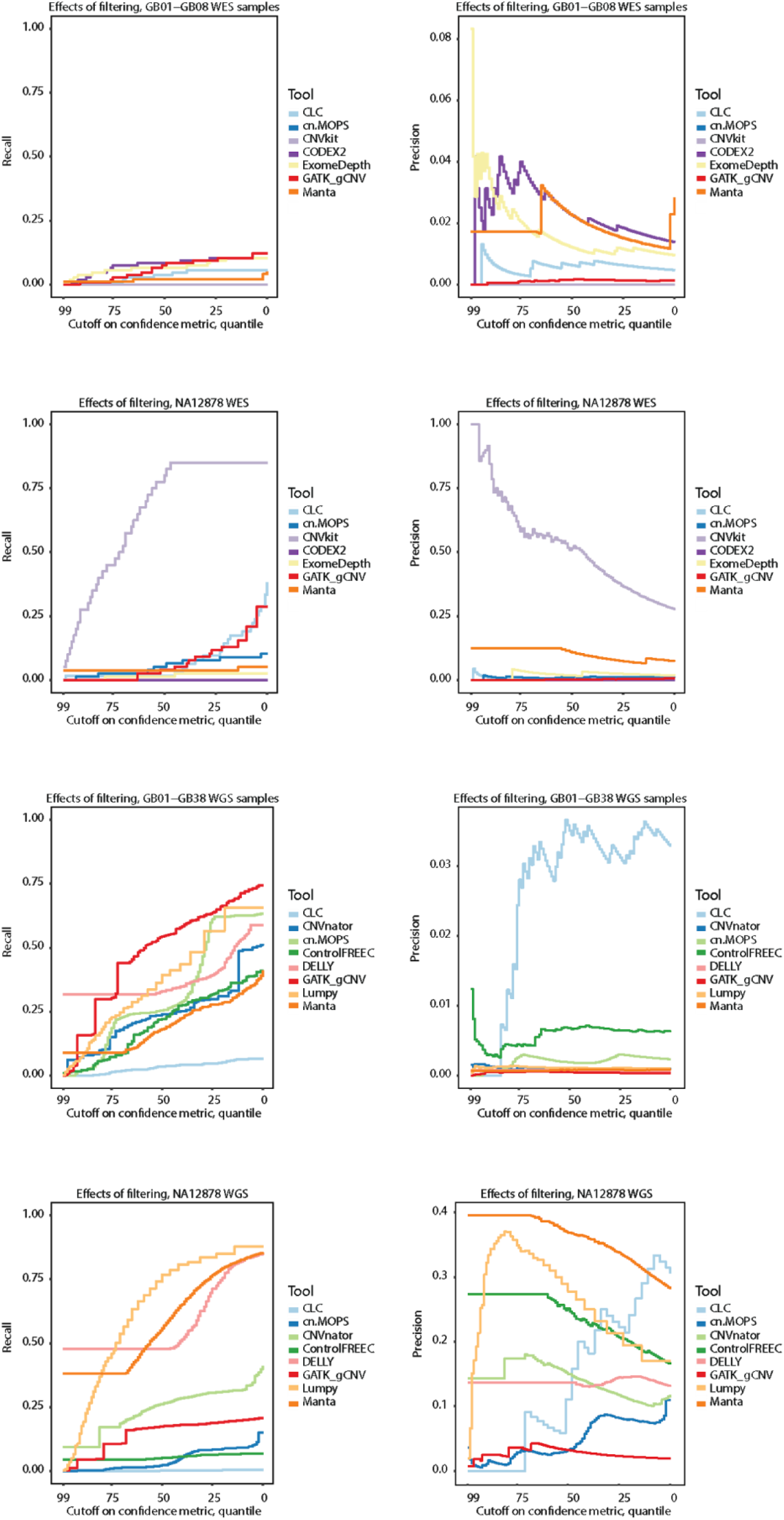
Recall and precision curves for GB01–08 and NA12878 WES samples, and GB01– GB38 and NA12878 WGS samples.

However, this performance of CNVkit was only seen for NA12878 sample and not for the remaining WES samples.

For WGS, recall for most tools decreased linearly as a function of filtering, the utility of which is therefore limited. Arguably, one exception for the GB01–GB38 WGS samples was GATK gCNV, which had proportionally good recall when filtering out the bottom 75^th^ percentile. However, such a cutoff did not improve the precision for the tool. For the WGS NA12878 sample, DELLY and Manta had good performance when selecting only the over-represented top confidence values, while Lumpy also displayed diminishing returns at lower scores. In the case of Manta, precision decreased predictably with filtering, while Lumpy had a somewhat unpredictable precision fraction curve, and DELLY’s precision appeared unaffected by filtering. CLC Genomics Workbench had repeated patterns for recall in both NA12878 and GB01–GB38 suggesting a relative ranking metric for each run. Collectively, sorting CNVs on confidence metrics from the tools did not offer any meaningful threshold for controlling precision, due to asymptotically positive recall curves (more liberal inclusion resulted in more hits). We further found several unpredictable precision curves, and overuse of the maximum confidence value, which gave a percentile of top-scoring CNVs without any additional metrics to rank. Exceptions to these conclusions were found only for the NA12878 sample.

### Short CNVs can be identified by NGS-based CNV calling tools

For the 38 WGS samples, DELLY and GATK gCNV called the most CNVs: 148,519 and 132,265, respectively. Manta, Lumpy, CNVnator, cn.MOPS, and Control-FREEC called 94,832, 93,166, 85,962, 36,491, and 13,160, respectively. CLC Genomics Workbench called 632 CNVs across the 29 samples it was run for. The tool was not run on all 38 samples due to computation time: it re-analyzes the base-level coverages of the control samples in every run, resulting in very long running times for WGS samples.

Tools with the same calling strategies had a higher overlap in called CNVs (Figure 5A). For example, DELLY, Lumpy, and Manta displayed a large degree of CNV overlap in WGS samples and they all use RP and SR information for calling CNVs. CNVnator, GATK gCNV, and DELLY called a high number of unique CNVs in WGS data which were not called by any other tool. Furthermore, a total 51.9% of all called CNVs were shorter than 1,000 bp. CNVs which were called by two or more tools were mostly short: less than 1,000 bp. Such CNVs are known to be less often called by array-based CNV calling approaches [8, 28]. Out of 407,671 CNVs called in the WGS samples, 74.4% were called only by a single tool. The percentages of CNVs called by 2–8 tools were 11.5, 9.5, 2.6, 0.8, 0.8, 0.4, and 0.1%, respectively (Figure 5A).

For WES (Figure 5B), CLC Genomics Workbench called 1,268 CNVs, cn.MOPS —787, CNVkit—66, CODEX2—762, ExomeDepth—1,123, GATK gCNV—7,116, and Manta— 174. The observed overlap between CNVs called between tools was lower than for WGS data: 90.3% of the total 9,944 CNVs were called by a single tool, and the percentages for 2–7 tools were 6.8, 2.0, 0.6, 0.2, 0.04, and 0%, respectively. GATK gCNV called the highest number of unique CNVs while the CNV calls by two or more tools were mostly longer than 1,000 bp. Despite the overlap between longer CNV calls, the majority (69.6%) of all CNV calls were shorter than 1,000 bp.

Lumpy, DELLY, Manta, and partly CNVnator performed best on NA12878 WGS data while CNVkit recalled almost all CNVs present in NA12878 WES dataset (Figure 5C–D). It is important to note that Manta and CNVnator were used for the generation of the NA12878 truth CNV set [8] and the CNV calls might have been favored to overfitting. More accurate picture of the tools’ performance can be obtained by evaluation of GB01–GB38 CNV calls.

Many CNVs confirmed by CytoScan HD SNP-array on WGS were called by multiple tools (Figure 5E). While 25.6% of all called CNVs overlapped in two or more tools (Figure 5A), more than 83.7% CNVs were overlapping with the CytoScan HD-confirmed CNV list (Figure 5E). As for WES, in more than two thirds of the cases where CytoScan identified a CNV, none of the tools called it. All tools performed similarly poorly on WES data (Figure 6) with cn.MOPS and CNVkit missing all the CNVs identified by CytoScan HD SNP-array.

**Figure 6.**
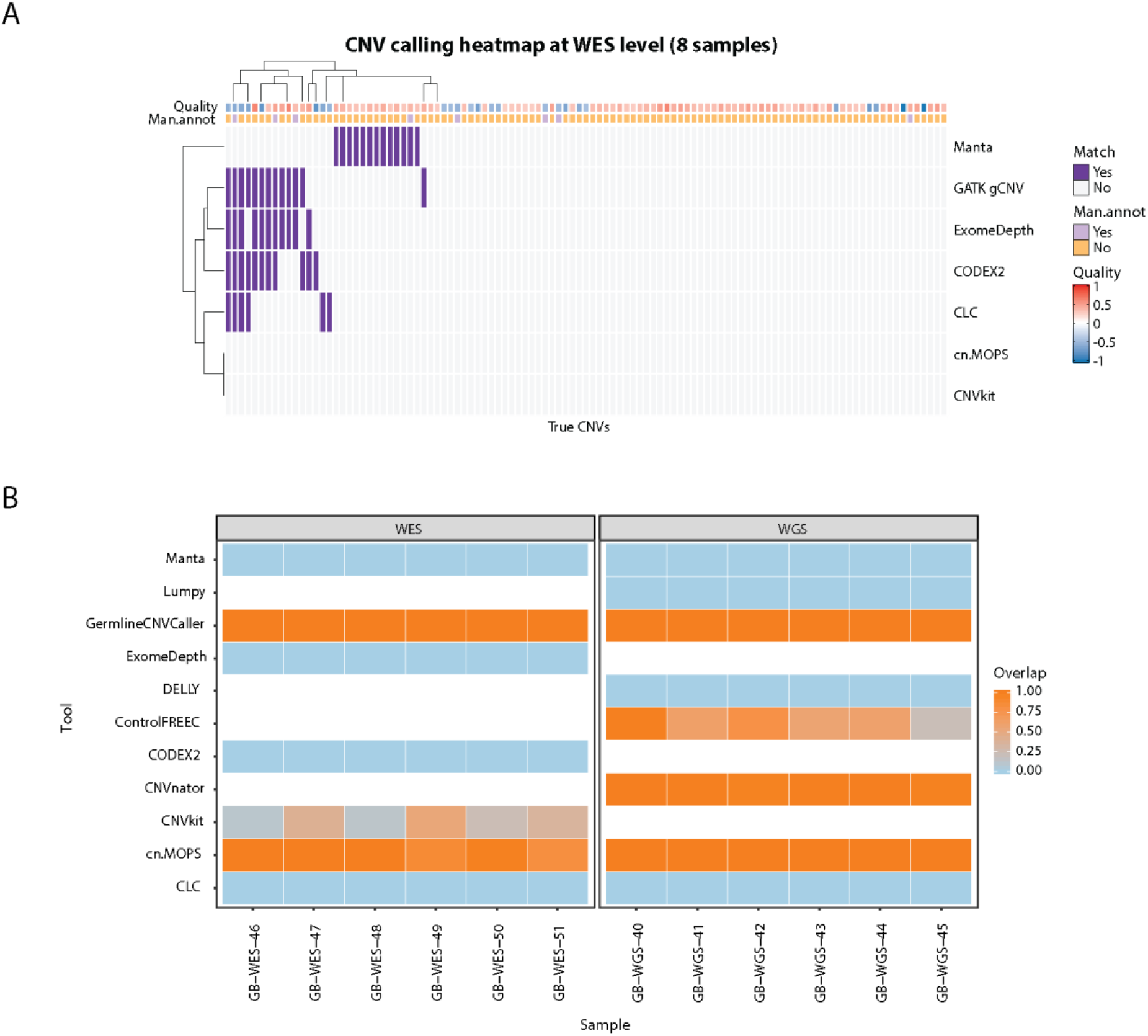
(A) CNV calling heatmap for 7 tools and 107 true CNVs at WES level in 8 samples (GB01–08). The Quality annotation represents the probe median score from CytoScan HD SNP-array and the Man.annot. refers to whether the CNV was independently manually confirmed. A positive quality score corresponds to duplications, and negative scores denote deletions. The order of rows/columns was determined using complete-linkage hierarchical clustering with Euclidean distance. (B) MLPA-confirmed CNV calling results for 11 CNV calling tools. GATK gCNV is labeled as GermlineCNVCaller.

### MLPA-confirmed CNV recall for CNV calling tools

To assess if tools accurately identify CNV breakends we used twelve MLPA-confirmed CNVs of varying sizes (1 exon to whole gene; deletions (N=11) or duplications (N=1), Supplementary Table 3) from six WES samples and six WGS samples (Figure 6B). Six out of 11 tools (CLC Genomics Workbench, CODEX2, DELLY, ExomeDepth, Lumpy, and Manta) did not identify any of the MLPA-confirmed CNVs. Conversely, GATK gCNV, CNVkit and cn.MOPS identified all MLPA-confirmed CNVs in WES. GAT K gCNV, cn.MOPS, CNVnator, and Control-FREEC identified all the CNVs confirmed by MLPA in WGS samples. CNVnator (WGS), cn.MOPS (WES), and CNVkit (WES) predicted shorter CNVs than the MLPA–defined truth, while GATK gCNV identified the full-length of both deleted and duplicated regions. Control-FREEC called all 6 CNVs in WGS samples but predicted shorter CNVs spanning 22.0–98.6% of true CNV length.

### Memory and CPU requirements for CNV calling tools

CPU and memory requirements were measured on a 28-core server grade cluster node, for the tools where it was possible to obtain an estimation on the NA12878 sample. Control-FREEC showed the best compromise between memory and CPU, both being low in WGS and even lower than requirements for other tools in WES (Figure 7). Memory-wise, DELLY and Manta were the other two tools with the lowest needs; the latter also having short computational times, while the former had one of the highest, possibly due to the fact that insertions and deletions were called subsequently and not in parallel. Surprisingly, cn.MOPS also showed low memory requirements on exomes, but the highest in genomes. However, it also offered one of the lowest computational times both in WES and WGS.

**Figure 7.**
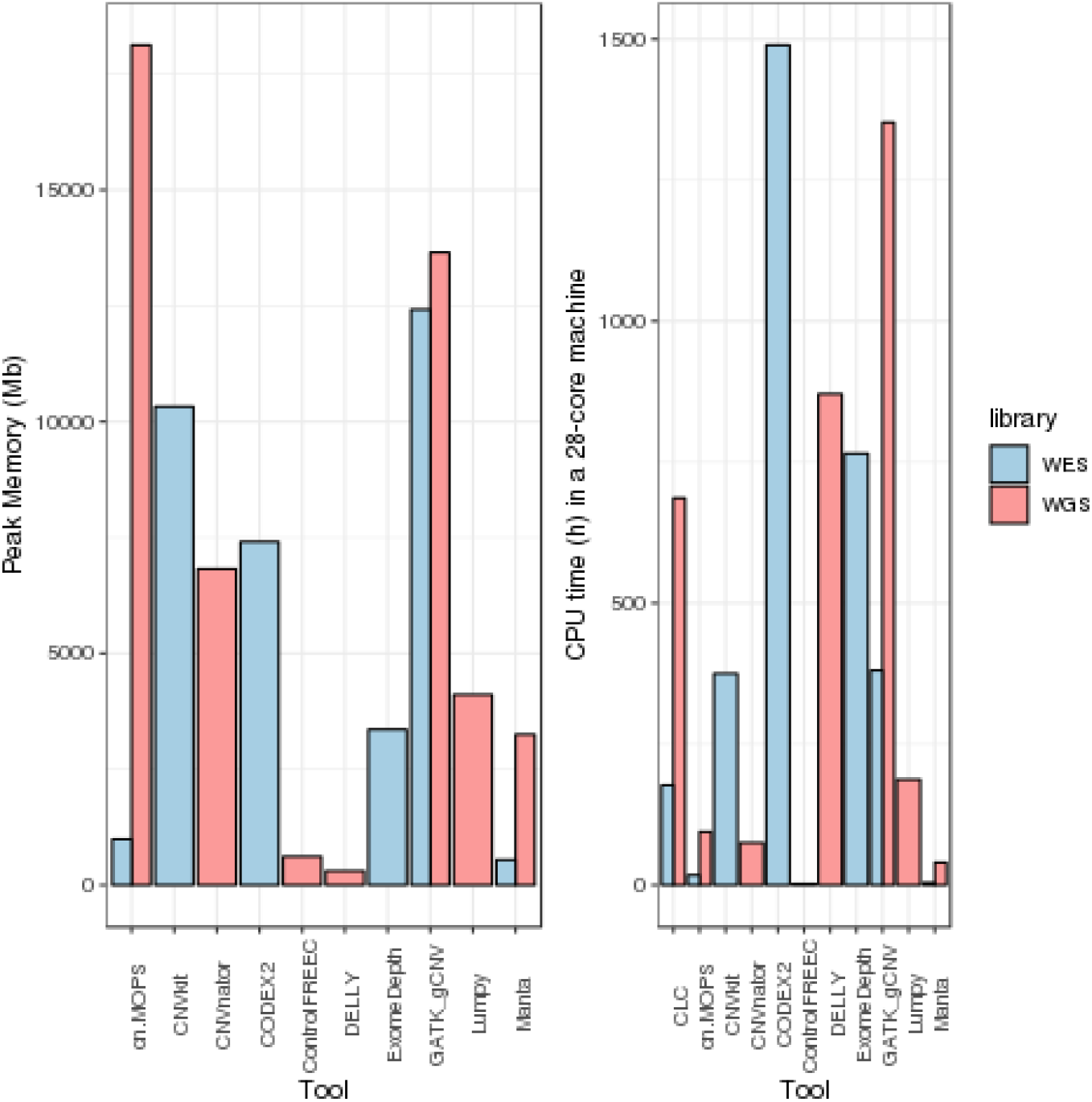
Maximum memory used by a tool measured in megabytes and total CPU time in hours run in 28-core machines with 128 GB RAM, while running NA12878. Some tools can distribute tasks over nodes, and total RAM usage is reported as total ma ximum.

GATK gCNV and cn.MOPS used a lot of RAM at peak memory, and it is possible that more RAM per node than the available 128GB per machine would have shortened the runtime by enabling better distribution of tasks. Computational requirements for creation of GATK’s model were not measured in this benchmark. Due to the batch caller nature of cn.MOPS, CNVkit, and CODEX2, many alignments have to be kept in memory at the same time, explaining the observed higher memory requirements.

## Discussion

From the reviewed 50 CNV calling tools, we observed that many of the tools were either not maintained with the last updates applied more than 5 years ago, or not widely used. We included 11 widely used or newly developed CNV calling tools, which fulfilled our selection criteria, to benchmark their performance on CNV calling on WES and WGS samples.

In order to establish a reference set of CNVs we used the best performing array-platform, in addition to NGS. CytoScan HD SNP-array technology was chosen as a sensitive and clinically adopted method to detect CNVs. It is important to note that because of the probe-dependent and genome-distributed nature of the array technology, not all short CNVs could be captured.

Therefore, the CNV calls classified as false positive (FP) in this benchmark should be interpreted carefully. Additionally, as CNV detection is a technically challenging task, none of the array-based standards in this study can ultimately be regarded as an absolute truth [8]. Besides the in-house GB01–GB38 samples, which were analyzed by Cytoscan HD SNP-array, we included the well-studied NA12878 sample, for which extensive efforts have been made to confirm all CNVs, based on several platforms, and NGS-based CNV callers. The latter might introduce a bias for these samples in this benchmark, as two of the included tools were used in this evaluation (Manta and CNVnator). Furthermore, the NA12878 sample and its truth CNV set are also popular for testing and optimising CNV tools, which could potentially explain the possible overfitting we observed e.g., for Manta and CNVkit, which had the highest discrepancy between NA12878 calls and calls on our cohort.

Tools with identical CNV calling strategy had a tendency to call the same CNVs, and, in general, read-depth based tools, or combinations including this strategy, performed best, when assessed on recall of CNVs from Cytoscan HP SNP-array, or NA12878 Gold Standard sample. The number of CNVs called varied more than a 100-fold; consequently, the recall rates for tools calling many CNVs were higher, and no systematic trade-off could be found to improve precision for these tools. In short, tools calling many CNVs hit the target more often, but high confidence CNVs were not generally showing a higher fraction of recall. For tools like DELLY and Lumpy, a combination of CNV metrics could be used to filter on the CNV calls as it is applied in SVTyper [29].

CNVs selected for experimental validation with MLPA were selected based on targeted gene panel sequencing, and were, therefore, not biased by CNV calls from tools tested in this analysis. It was, however, striking that tools could be split into two groups: those that were able to recall all six independent CNVs and those that called none.

GATK gCNV caller performed best at CNV recall and is clearly the most sensitive tool for CNV identification for both WES and WGS data, but comes with poor precision, like all tools tested (highest precision mean < 13% for WES and < 6% for WGS). GATK gCNV is also the best performing tool when recalling MLPA-confirmed CNVs and estimating their breakends correctly, even if four other tools also recalled the CNVs. The good performance of GATK gCNV and cn.MOPS caller comes at a high computational cost and the former was almost twice as computationally expensive as the third-highest consuming WGS tool considering CPU/h and peak RAM usage.

The GnomAD database [14] shows how CNV calls can be used clinically, but more research and larger cohort studies are needed for better annotation and inference of causation of CNVs. Our study shows that more work has to be done on collecting large and well-annotated datasets with CNV detection on several platforms, in order to drive the development of tools with improved precision on CNV calling from NGS data. The current state of tools for finding CNVs is suited for identifying complex traits in large cohorts, for which we suggest to use the overlap between several tools. Using rare CNVs called from NGS as a basis for genome-wide association studies is not currently advisable.

The future for NGS-based CNV calling tools is likely to rely on the utilization of a combination of long- and short-read sequencing [30]. This is particularly true considering the need for CNV annotation that explains causative traits and which will require sequencing of large cohorts with two simultaneous protocols. Alternatively, future improvements on both price and error rate for long-read sequencing are needed. In terms of filtering CNV calling results, better annotation of CNVs is a clear optimization candidate. The ability to leverage additional genetic data, such as RNA-seq, or even static knowledge of genetic sites from associations to epigenetic mechanisms or regulation, may also guide the selection and prioritisation process in the near future. In a clinical setting, production of background panels or databases to filter true common CNVs or common FPs called by each tool can greatly reduce the number of relevant CNVs presented for interpretation (data not shown), just like databases like gnomAD SV [31] can be used to reduce the numbers of common CNVs. Lastly, transcriptional regulation of the altered regions requires more investigations, so that the causative effect of CNVs can be elucidated, and potentially be predicted in each case.

Our work has several limitations. First, we benchmarked only a limited set of tools; however, findings are in line with larger studies [32], relying on single truth sets. Furthermore, the observed potential for overfitting to NA12878 Gold Standard sample by some tools complicated the accurate evaluation of recall and precision with a well-annotated dataset. Finally, the main limitation of our work is the lack of well-defined true CNV sets, therefore our analysis using CytoScan HD SNP-array calls vastly underestimates CNV call precision on the in-house data sets, but this caveat should not favor specific tools.

In summary, by reviewing 50 tools for CNV calling, of which 11 were included for a benchmark (CLC Genomics Workbench (WGS and WES), cn.MOPS (WGS and WES), CNVkit (WES), CNVnator (WGS), CODEX2 (WGS), Control-FREEC (WGS), DELLY (WGS), ExomeDepth (WES), GATK gCNV (WES and WGS), Lumpy (WGS), and Manta (WES and WGS)), we conclude that CNV identification from NGS data remains challenging. For the best reliability of CNV calling from NGS data, we observed that even if the tools were developed for WES data or allowed it as input, they did not perform well. we suggest WGS as the only NGS-based option for broad calling of CNVs. Furthermore, low precision in all tools leads us to recommend a hypothesis-based approach for finding causative CNVs by NGS in the clinic, and further validation of these candidates by manual inspection, MLPA or array - based approaches. If multiple samples are available from the same protocol, we suggest using these to filter by commonly called CNVs. If only the WGS data is available for the sample, for a higher precision of CNV calls, multiple CNV calling tools should be used. we suggest combining tools which have the best recall: GATK gCNV, Lumpy, DELLY, and cn.MOPS.

## Methods

### Sequencing and read alignments

Genomic DNA (gDNA) was extracted from whole blood samples using a liquid handling automated station (Tecan). WES was performed from 200 ng of gDNA. Fragmentation was done on Covaris S2 (Agilent) to approximately 300 base pair fragments and adaptor ligation was performed using KAPA HTP Library Preparation Kit. Exomes were enriched with SureSelectXT Clinical Research Exome kit (Agilent). Paired-end sequencing with average read depth of at least 50x was performed using HiSeq2500 or NextSeq500 platforms from Illumina. For WGS analysis, sequencing libraries were prepared from 500 ng gDNA using Nextera DNA Flex library prep kit (Illumina), according to the manufacturer’s instructions. WGS libraries were sequenced on Illumina NovaSeq6000 with sequencing depth of at least 30x. Sequenced reads were trimmed and aligned to the human reference genome (hg19/GRCh37) using BWA MEM 0.7.12 software [33].

### Selection of Normals

CNVkit, CODEX2, ExomeDepth, and GATK gCNV require a group of samples to represent healthy control genomes. Normal samples should be produced with the same technical protocol, thus presenting a similar pattern of technical noise. Due to the nature of our experimental set up, no healthy control samples were available to create the Panel of Normals. Normal-like samples were selected instead. Seventy whole genomes were chosen as WGS normals. Ninety-four exome samples were chosen as WES normals.

### MLPA

MLPA analysis was performed according to the manufacturer’s instructions (MRC-Holland, Amsterdam, the Netherlands) using appropriate MLPA Kits for BRCA1 (NM_00007294), BRCA2 (NM_000059), FLCN (NM_144997), MSH2 (NM_000251), MSH6 (NM_000179), PALB2 (NM_024675), PMS2 (NM_000535), VHL (NM_000551).

### CytoScan HD SNP-array

gDNA was isolated using the liquid handling automated station (Tecan). Purified DNA was quantified using the Qubit instrument (Life Technologies). CytoScan HD SNP-array (Thermofisher Scientific), which contains 2.67 million genome-wide markers, was performed on extracted DNA from samples GB01–GB38, according to the manufacturer’s instructions. Result files were analysed using Nexus Copy Number software 10.0 (BioDiscovery) using NCBI Build 37 as reference. The samples were pre-processed by systematic correction (Quadratic), probes were recentered by the median and applying the mean of Combine Replicates Between Arrays. Subsequently, data were processed by SNP-FASST2 Segmentation with a significance threshold of 1.0E-8 and max contiguous probe spacing of 1000 Kbp with a minimum of 3 probes per segment. Following thresholds were applied for calling CNVs: High Gain = 0.7; Gain = 0.23; Loss =-0.37; Big Loss =-1.1 and heterozygous imbalance threshold of 0.4. All gains or losses not covered by an allelic imbalance event were considered as false-positive and removed. Moreover, independently of the automatically generated CNV calls, each sample was visually inspected for CNVs using Nexus Copy Number software 10.0 (BioDiscovery).

### NA12878 Gold Standard

Gold standard for NA12878 CNVs was produced by the 1000 Genomes Project. It contains only high confidence CNVs and the list of all CNVs was obtained from Haraksingh *et al*., 2017 [8]. In total, 2,076 CNVs of 51–453,313 bp sizes were used for CNV calling software evaluation. We have performed NA12878 WGS sequencing in-house while WES data was obtained from: ftp-trace.ncbi.nih.gov/giab/ftp/data/NA12878/Garvan_NA12878_HG001_HiSeq_Exome/.

### CNVnator

CNVnator [17] was run using the default parameters and recommendations from the authors with the bin size of 100 for all WGS samples.

### CLC Genomics Benchmark

CLC Genomics Benchmark uses fastq files as input to the CNV calling workflow which are subsequently mapped to the reference genome by their internal read mapping tool [34]. The mapped reads of the samples under investigation and the control samples are then used for calling CNVs. The called CNVs can be exported as BED files.

### GATK gCNV

GATK gCNV [19] calling is composed of two workflows: model creation and individual sample calling.

During model creation, the Panel of Normals of WGS and WES composed by 70 and 94 samples respectively, were generated. For WGS, intervals of 1000 bp and 0 bp padding were produced and filtered following GATK recommendations. No intervals were generated for WES samples. Read counts were measured in the exome regions and whole genome intervals and ploidy models were generated. Finally, a model per chromosome was produced using GermlineVariantCall. Benchmark samples were subsequently run, following the same procedure as the Panel of Normals, but including the available models during ploidy determination and germline variant call. Chromosomal calls were finally merged using PostprocessGermlineCNVCalls.

### DELLY

Duplications and deletions were called by using DELLY [25]. Only one library size was available and it was provided at a time. Each of the samples only contained one read-group and no further specifications were given. The variant call was performed searching for duplication events (-t DUP) and deletions (-t DEL). The resulting bcf files were merged using bcftools concat.

### cn.MOPS

cn.MOPS [20] was used to call CNVs on all WES and WGS samples following the vignette [35]. Due to memory constraints, WGS sample bam files were grouped together in batches of 11 to 12 samples before analysis, and each batch was run as follows. Using the provided getReadCountsFromBAM function, read counts were extracted from the bam files for chromosomes 1–22 using a window length parameter WL=500 in order to achieve roughly 50– 100 reads per window. The cn.mops function was used to call CNVs and integer copy numbers were extracted using calcIntegerCopyNumbers.

For the WES samples, all bam files were run together as a single batch. A bed file with the targeted regions was converted into GRanges format and used to extract read counts in the regions with the getSegmentReadCountsFromBAM function. CNVs were then called using the exome cn.mops function and integer copy numbers calculated as above.

### CNVkit

CNVkit [22] was used to call CNVs on all WES and WGS samples, grouped together by library type and according to the authors guidelines [36]. Briefly, we used the default parameters, restricting the analyses to the WGS mappable regions and, in the case of WES, to the captured regions also.

### Control-FREEC

Control-FREEC 11.5 [24, 37] was used to process the 39 WGS samples of the benchmark starting from BAM files. Default parameter values (forceGCcontentNormalization = 0, minCNAlength = 1, coefficientOfVariation = 0.05) were used, however, ploidy was set to 2, mateOrientation to “FR” for Illumina paired-end reads, and the sex of each sample was supplied (two males and seven females). In this case, the hg19 reference used for alignment was used and mappability of the genome was disregarded (read length = 151). Sambamba 0.6.7 [38], BEDTools 2.27.1 [39], Samtools 1.9 [40, 41], and R 3.5.0 [42] were used as dependencies. The script ‘assess_significance.R’ provided by the Control-FREEC developers was used to add p-values to the detected CNVs.

### Manta

Manta [26] was used to infer deletions and duplications from both WES and WGS sequence data using default parameters.

### LUMPY

For each of our WGS sample bam files, we used the LUMPY Express wrapper [27] to call CNVs as described in the official documentation on GitHub (https://github.com/arq5x/lumpy-sv): discordants were extracted with samtools view, filtering out reads by flag 1294; split reads were extracted using the provided extractSplitReads_BwaMem script. Both were sorted and then provided as input to the lumpyexpress utility together with the original bam to output a vcf giving structural variants, from which we obtained CNVs as the called deletions and duplications.

### ExomeDepth

ExomeDepth [21] was used to detect CNVs from WES sequence data following the authors recommended best practices [43]. Since ExomeDepth takes advantage of the tight correlation structures between large numbers of samples when building a reference sample, we used our reference samples as a Panel of Normals. Even though these reference samples are not ideal (since they are not from the same batch), they are a good representation of the samples in our laboratory, as well as the experimental design most users will encounter (related samples from the same lab, but not the same batch).

### CODEX2

To run CODEX2 [23], we grouped all the WES samples together with 50 extra Panel of Normals WES samples. This batch of samples was then analysed for each of chromosomes 1 through 22 by following the provided documentation from GitHub (https://github.com/yuchaojiang/CODEX2), using default or suggested parameters and functions except where otherwise noted. We used CODEX2 without specifying negative control samples, using the normalize_null function for normalization. For chromosomes 1, 4, 6, and 14, the glm.fit procedure in CODEX2 did not converge, despite increasing the parameter K = 1:10 as per the authors’ suggestion.

### Data processing and plotting

All tools were run using Snakemake [44], post-processing of the called CNVs was carried out using Python 3.6 and R 3.5.0 [42]. Scripts for running the tools are available on GitHub (https://github.com/cphgeno/CNVbench). The rtracklayer package [45] was used for main processing of the files, and ggplot2 [46], ComplexHeatmap [47], and RColorBrewer [48] were the key resources for plotting.

### Fraction curve generation

The analyzed data was split into four data sets: NA12878 WES, GB01–GB08 WES, NA12878 WGS, and GB01–GB38 WES. For each data set, and for each tool that called CNVs for the dataset in question, we filtered the CNV calls according to a confidence metric provided by the tool itself (see below). Specifically, WESubset the tool data set on every confidence metric percentile from 0 to 99 by taking all CNV calls with confidence scores less/greater (as appropriate for the metric) than or equal to a given percentile. Recall and precision were then calculated and plotted for the resulting, filtered CNV calls.

Note that some tools use only a few discrete confidence values or use a single value for a large proportion of the calls. Therefore, a cutoff at the 99th percentile does not necessarily contain 1% of the called CNVs if, for example, 50% of the CNVs in the call set are assigned the best confidence score.

The confidence metrics used for each tool were as follows. CLC Genomics Workbench: Absolute fold change; CNVkit: Mean squared standard error of log2 of the copy number; CNVnator: t-statistic p-value; cn.MOPS: Median informative/non-informative ratio value; CODEX2: Likelihood ratio; ControlFREEC: Wilcoxon rank sum test p-value; DELLY: Genotype quality values; ExomeDepth: Observed/expected read ratio; GATK gCNV: CNQ scores (difference between the two best genotype Phred-scaled log posteriors); Lumpy: Number of pieces of evidence supporting the variant across all samples; Manta: CNV quality score.

### Performance profiling

Snakemake’s built-in capabilities for benchmarking runtime and memory usage were used to measure wall-clock time and peak resident set size for calling CNVs on sample NA12878. Tools were tested on a HP Apollo 6000 System ProLiant XL230a Gen9 Server blade, on a node with 28 64-bit Intel Xeon E5-2683 v3 @2.00 GHz CPUs available, and 128 GB, DDR4 @2133 MHz RAM.

## List of abbreviations

AS: Assembly
CNV: Copy number variation
gDNA: genomic DNA
MLPA: Multiplex ligation-dependent probe amplification
NGS: Next generation sequencing
RD: Read-depth
RP: Read-pair
SR: Split-read
SV: Structural variant
WES: Whole exome sequencing
WGS: Whole genome sequencing

## Declarations

### Availability of data and material

WGS data for NA12828 and snakemake workflow to run and test all tools in this benchmark is deposited at: https://github.com/cphgeno/CNVbench.

WES for NA12828 WES is from: ftp://ftp-trace.ncbi.nih.gov/giab/ftp/data/NA12878/Garvan_NA12878_HG001_HiSeq_Exome

### Ethics approval and consent to participate

DNA samples originate from the in-house anonymized quality samples.

## Acknowledgements

The authors would like to thank Denise Serra for editing and typesetting of figures.

## Author contributions

FOB conceived and designed the study. MG, MHT, MSR, FGV, CBP, SAS, and SK tested tools, processed the output, and developed the computational framework. MBM, CWY, OØ, and MR gathered the experimental data and performed analysis of the CytoScan HD SNP - arrays. RLM, LRO, FCN, OW, and FOB supervised the work. All authors discussed the results. MG, MHT, MSR, FGV, CBP, SAS, SK, MBM, CWY, LRO, and FOB contributed to the final manuscript.

## Bibliography

1. Ionita-Laza I, Rogers AJ, Lange C, Raby BA, Lee C. Genetic association analysis of copy - number variation (CNV) in human disease pathogenesis. Genomics. 2009;93:22–6. doi:10.1016/j.ygeno.2008.08.012.

2. Sebat J, Lakshmi B, Troge J, Alexander J, Young J, Lundin P, et al. Large-scale copy number polymorphism in the human genome. Science. 2004;305:525–8. doi:10.1126/science.1098918.

3. Takumi T, Tamada K. CNV biology in neurodevelopmental disorders. Curr Opin Neurobiol. 2018;48:183–92. doi:10.1016/j.conb.2017.12.004.

4. Kumaran M, Cass CE, Graham K, Mackey JR, Hubaux R, Lam W, et al. Germline copy number variations are associated with breast cancer risk and prognosis. Sci Rep. 2017;7:14621. doi:10.1038/s41598-017-14799-7.

5. Iafrate AJ, Feuk L, Rivera M N, Listewnik ML, Donahoe PK, Qi Y, et al. Detection of large-scale variation in the human genome. Nat Genet. 2004;36:949–51. doi:10.1038/ng1416.

6. Zarrei M, MacDonald JR, Merico D, Scherer SW. A copy number variation map of the human genome. Nat Rev Genet. 2015;16:172–83. doi:10.1038/nrg3871.

7. Nowakowska B. Clinical interpretation of copy number variants in the human genome. J Appl Genet. 2017;58:449–57. doi:10.1007/s13353-017-0407-4.

8. Haraksingh RR, Abyzov A, Urban AE. Comprehensive performance comparison of high - resolution array platforms for genome-wide Copy Number Variation (CNV) analysis in humans. BMC Genomics. 2017;18:321. doi:10.1186/s12864-017-3658-x.

9. Zhao M, Wang Q, Wang Q, Jia P, Zhao Z. Computational tools for copy number variation (CNV) detection using next-generation sequencing data: features and perspectives. BMC Bioinformatics. 2013;14 Suppl 11:S1. doi:10.1186/1471-2105-14-S11-S1.

10. Alkan C, Coe BP, Eichler EE. Genome structural variation discovery and genotyping. Nat Rev Genet. 2011;12:363–76. doi:10.1038/nrg2958.

11. Pirooznia M, Goes FS, Zandi PP. Whole-genome CNV analysis: advances in computational approaches. Front Genet. 2015;6:138. doi:10.3389/fgene.2015.00138.

12. Kent WJ, Sugnet CW, Furey TS, Roskin KM, Pringle TH, Zahler AM, et al. The human genome browser at UCSC. Genome Res. 2002;12:996–1006. doi:10.1101/gr.229102.

13. Kaminsky EB, Kaul V, Paschall J, Church DM, Bunke B, Kunig D, et al. An evidence - based approach to establish the functional and clinical significance of copy number variants in intellectual and developmental disabilities. Genet Med. 2011;13:777–84. doi:10.1097/GIM.0b013e31822c79f9.

14. Karczewski KJ, Francioli LC, Tiao G, Cummings BB, Alföldi J, Wang Q, et al. Variation across 141,456 human exomes and genomes reveals the spectrum of loss-of-function intolerance across human protein-coding genes. BioRxiv. 2019. doi:10.1101/531210.

15. Yao R, Zhang C, Yu T, Li N, Hu X, Wang X, et al. Evaluation of three read-depth based CNV detection tools using whole-exome sequencing data. Mol Cytogenet. 2017;10:30. doi:10.1186/s13039-017-0333-5.

16. Scionti F, Di Martino MT, Pensabene L, Bruni V, Concolino D. The cytoscan HD array in the diagnosis of neurodevelopmental disorders. High-Throughput. 2018;7. doi:10.3390/ht7030028.

17. Abyzov A, Urban AE, Snyder M, Gerstein M. CNVnator: an approach to discover, genotype, and characterize typical and atypical CNVs from family and population genome sequencing. Genome Res. 2011;21:974–84. doi:10.1101/gr.114876.110.

18. Zhao Q, Caballero OL, Levy S, Stevenson BJ, Iseli C, de Souza SJ, et al. Transcriptome - guided characterization of genomic rearrangements in a breast cancer cell line. Proc Natl Acad Sci USA. 2009;106:1886–91. doi:10.1073/pnas.0812945106.

19. Babadi M, Lee SK, Smirnov A, Lichtenstein L, Gauthier LD, Howrigan DP, et al. Abstract 2287: Precise common and rare germline CNV calling with GATK. Cancer Res. 2018;78 13 Supplement:2287–2287. doi:10.1158/1538-7445.AM2018-2287.

20. Klambauer G, Schwarzbauer K, Mayr A, Clevert D -A, Mitterecker A, Bodenhofer U, et al. cn.MOPS: mixture of Poissons for discovering copy number variations in next-generation sequencing data with a low false discovery rate. Nucleic Acids Res. 2012;40:e69. doi:10.1093/nar/gks003.

21. Plagnol V, Curtis J, Epstein M, Mok KY, Stebbings E, Grigoriadou S, et al. A robust model for read count data in exome sequencing experiments and implications for copy number variant calling. Bioinformatics. 2012;28:2747–54. doi:10.1093/bioinformatics/bts526.

22. Talevich E, Shain AH, Botton T, Bastian BC. CNVkit: Genome-Wide Copy Number Detection and Visualization from Targeted DNA Sequencing. PLoS Comput Biol. 2016;12:e1004873. doi:10.1371/journal.pcbi.1004873.

23. Jiang Y, Wang R, Urrutia E, Anastopoulos IN, Nathanson KL, Zhang NR. CODEX2: full-spectrum copy number variation detection by high-throughput DNA sequencing. Genome Biol. 2018;19:202. doi:10.1186/s13059-018-1578-y.

24. Boeva V, Popova T, Bleakley K, Chiche P, Cappo J, Schleiermacher G, et al. Control - FREEC: a tool for assessing copy number and allelic content using next-generation sequencing data. Bioinformatics. 2012;28:423–5. doi:10.1093/bioinformatics/btr670.

25. Rausch T, Zichner T, Schlattl A, Stütz AM, Benes V, Korbel JO. DELLY: structural variant discovery by integrated paired-end and split-read analysis. Bioinformatics. 2012;28:i333–9. doi:10.1093/bioinformatics/bts378.

26. Chen X, Schulz-Trieglaff O, Shaw R, Barnes B, Schlesinger F, Källberg M, et al. Manta: rapid detection of structural variants and indels for germline and cancer sequencing applications. Bioinformatics. 2016;32:1220–2. doi:10.1093/bioinformatics/btv710.

27. Layer RM, Chiang C, Quinlan AR, Hall IM. LUMPY: a probabilistic framework for structural variant discovery. Genome Biol. 2014;15:R84. doi:10.1186/gb-2014-15-6-r84.

28. Carter NP. Methods and strategies for analyzing copy number variation using DNA microarrays. Nat Genet. 2007;39 7 Suppl:S16–21. doi:10.1038/ng2028.

29. Chiang C, Layer RM, Faust GG, Lindberg MR, Rose DB, Garrison EP, et al. SpeedSeq: ultra-fast personal genome analysis and interpretation. Nat Methods. 2015;12:966–8. doi:10.1038/nmeth.3505.

30. Sanchis-Juan A, Stephens J, French CE, Gleadall N, Mégy K, Penkett C, et al. Complex structural variants in Mendelian disorders: identification and breakpoint resolution using short- and long-read genome sequencing. Genome Med. 2018;10:95. doi:10.1186/s13073-018-0606-6.

31. Collins RL, Brand H, Karczewski KJ, Zhao X, Alföldi J, Francioli LC, et al. A structural variation reference for medical and population genetics. Nature. 2020;581:444–51. doi:10.1038/s41586-020-2287-8.

32. Kosugi S, Momozawa Y, Liu X, Terao C, Kubo M, Kamatani Y. Comprehensive evaluation of structural variation detection algorithms for whole genome sequencing. Genome Biol. 2019;20:117. doi:10.1186/s13059-019-1720-5.

33. Li H. Aligning sequence reads, clone sequences and assembly contigs with BWA-MEM. arXiv preprint 13033997. 2013.

34. QIAGEN. White paper on CLC read mapper [White paper]. 2012. http://resources.qiagenbioinformatics.com/white-papers/White_paper_on_CLC_read_mapper.pdf. Accessed 24 Apr 2019.

35. Klambauer G. cn.mops - Mixture of Poissons for CNV detection in NGS data. Software Manual. 2019;1.30.0.

36. Talevich E. Copy number calling pipeline. CNVkit. https://cnvkit.readthedocs.io/en/stable/pipeline.html. Accessed 14 May 2019.

37. Boeva V, Zinovyev A, Bleakley K, Vert J -P, Janoueix-Lerosey I, Delattre O, et al. Control-free calling of copy number alterations in deep-sequencing data using GC-content normalization. Bioinformatics. 2011;27:268–9. doi:10.1093/bioinformatics/btq635.

38. Tarasov A, Vilella AJ, Cuppen E, Nijman IJ, Prins P. Sambamba: fast processing of NGS alignment formats. Bioinformatics. 2015;31:2032–4. doi:10.1093/bioinformatics/btv098.

39. Quinlan AR, Hall IM. BEDTools: a flexible suite of utilities for comparing genomic features. Bioinformatics. 2010;26:841–2. doi:10.1093/bioinformatics/btq033.

40. Li H, Handsaker B, Wysoker A, Fennell T, Ruan J, Homer N, et al. The Sequence Alignment/Map format and SAMtools. Bioinformatics. 2009;25:2078–9. doi:10.1093/bioinformatics/btp352.

41. Li H. A statistical framework for SNP calling, mutation discovery, association mapping and population genetical parameter estimation from sequencing data. Bioinformatics. 2011;27:2987–93. doi:10.1093/bioinformatics/btr509.

42. R Core Team. R: A Language and Environment for Statistical Computing. 2020.

43. Plagnol V. ExomeDepth Vignette. 2016;1.1.10.

44. Köster J, Rahmann S. Snakemake--a scalable bioinformatics workflow engine. Bioinformatics. 2012;28:2520–2. doi:10.1093/bioinformatics/bts480.

45. Lawrence M, Gentleman R, Carey V. rtracklayer: an R package for interfacing with genome browsers. Bioinformatics. 2009;25:1841–2. doi:10.1093/bioinformatics/btp328.

46. Wickham H. ggplot2: Elegant Graphics For Data Analysis. Springer-Verlag New York; 2016.

47. Gu Z, Eils R, Schlesner M. Complex heatmaps reveal patterns and correlations in multidimensional genomic data. Bioinformatics. 2016;32:2847–9. doi:10.1093/bioinformatics/btw313.

48. Neuwirth E. RColorBrewer: ColorBrewer Palettes. 2014.

