## Supplementary figures and images for "A comparison of tools for copy-number variation detection in germline whole exome and whole genome sequencing data"

### Supplementary Figure 1

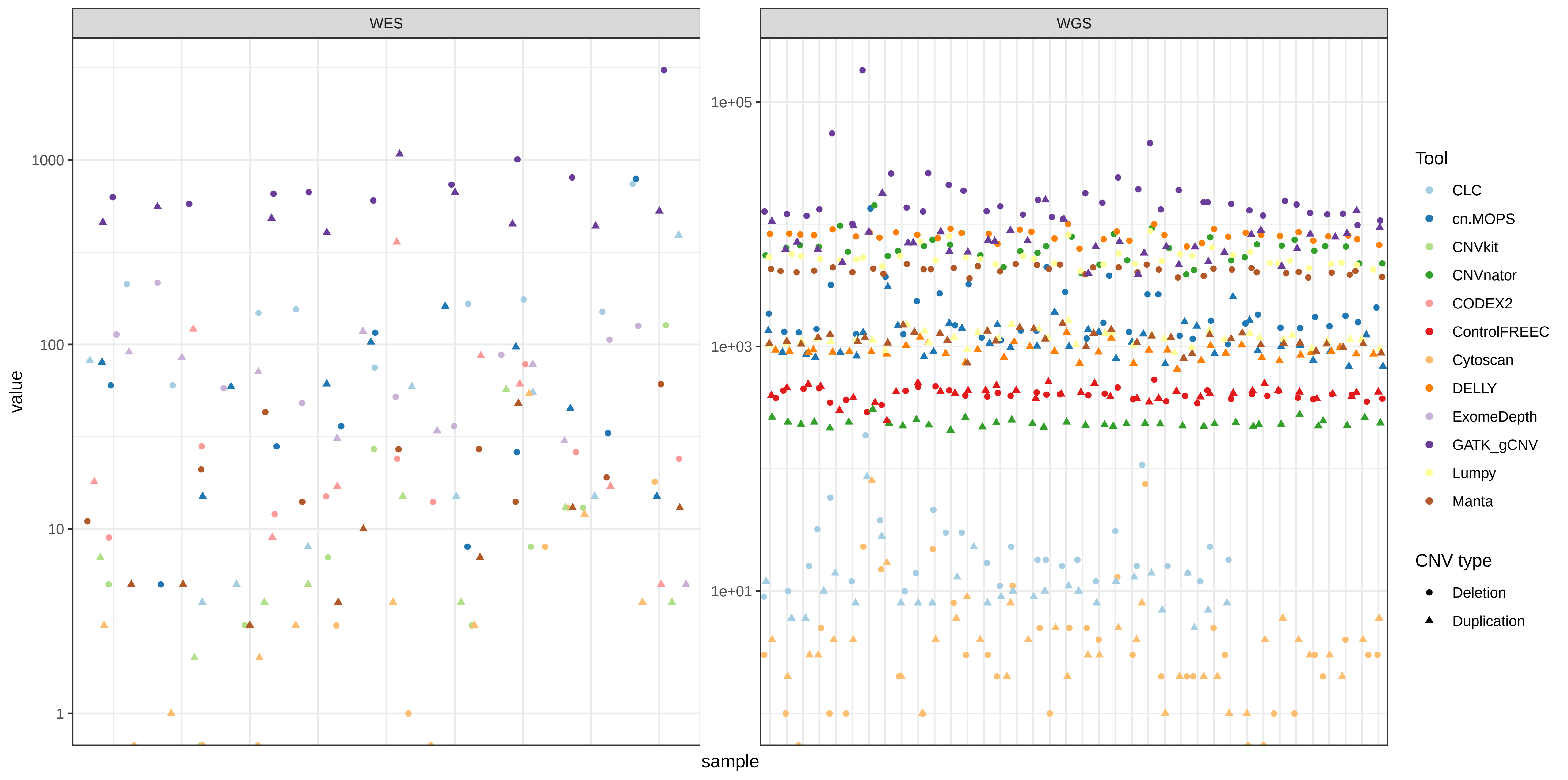

### Supplementary Figure 2

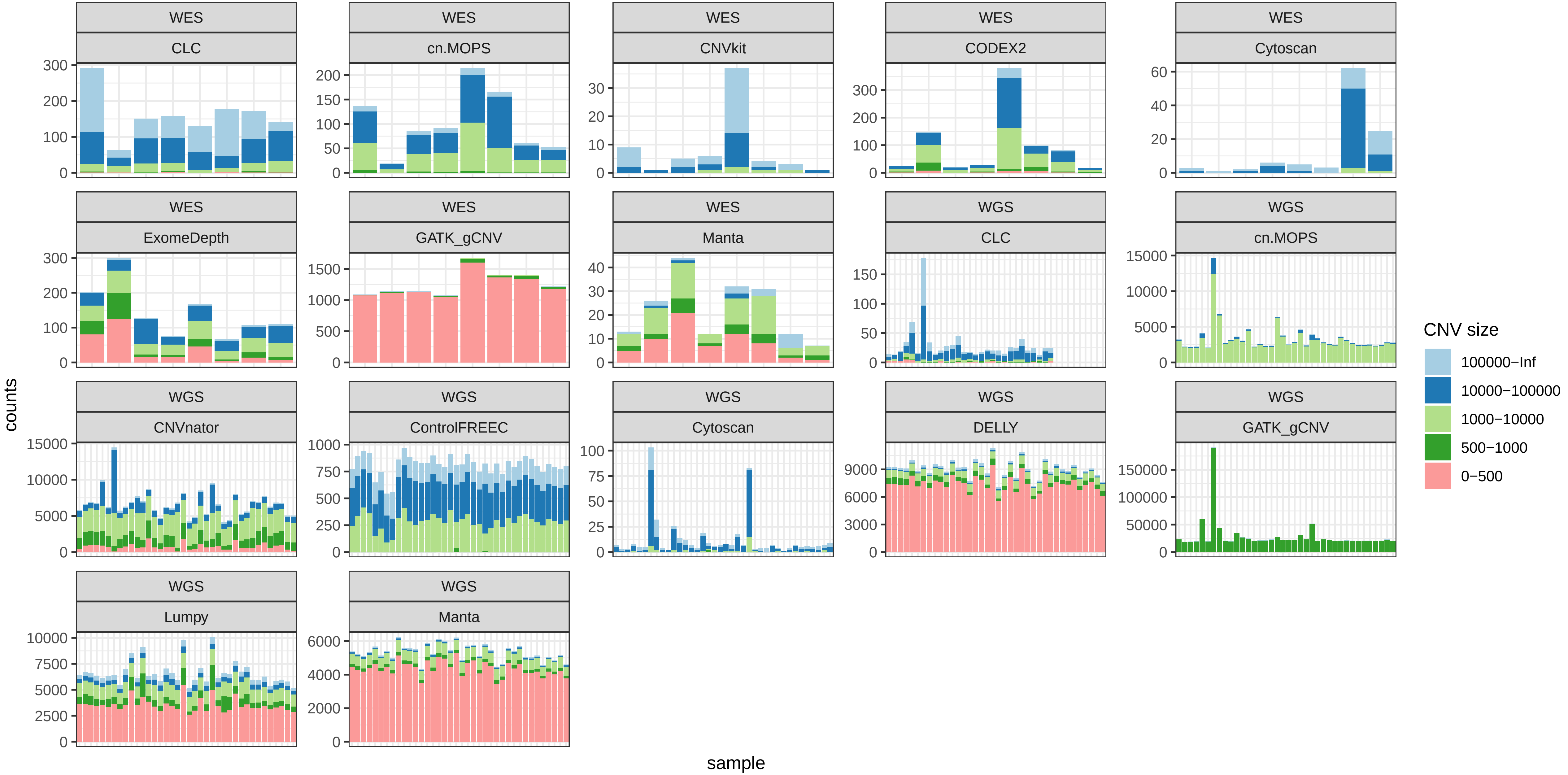
