## Supplementary File 1 for "A comparison of tools for copy-number variation detection in germline whole exome and whole genome sequencing data"

**CNVnator**

CNVnator tool discovers CNVs from RD analysis of WGS data. The method is based on combination of mean-shift, multiple-bandwidth partitioning and GC correction approaches. CNVnator was calibrated using 1000 Genomes Project and enables atypical CNV detection.

**CLC Genomics Benchmark**

CLC Genomics Benchmark uses a RD method for CNV detection in WES and WGS samples. A coverage confidence region is determined by the control samples. Any regions with unexpectedly low or high coverage are defined as CNVs. This is done by setting up a statistical model for the variation in fold-change as a function of coverage in the baseline. If the fold-change in specific regions are significantly different than the expected one, then these regions are called as CNVs [[26]](http://f1000.com/work/citation?ids=6951269&pre=&suf=&sa=0). The method is inspired by the studies of Li et al, 2012 [[27]](http://f1000.com/work/citation?ids=2016131&pre=&suf=&sa=0) and Niu and Zhang, 2012 [[28]](http://f1000.com/work/citation?ids=5057684&pre=&suf=&sa=0).

**GATK gCNV**

GATK’s pipeline for producing CNV germline calls is based on RD. Its algorithm is able to identify batch and in-sample depth biases and simultaneously infer copy-number estates. The pipeline consists of two different parts. First, a group of samples from the same batch are used for generating a ploidy model, which determines baseline contig ploidies, and a copy-number denoising model (cohort model). Such models are later used for determining the germline ploidy baseline level and CNV calls on individual samples (case model). The pipeline is able to model and determine copy number biases and variants in both WGS and WES.

**DELLY**

DELLY is an SV calling software which is able to combine short insert read-pairs, long insert mate-pairs and SR information to identify copy number (tandem and deletions) among other variants. In this benchmark, only short insert paired-end reads were used. DELLY’s algorithm is based on discordant pairs identification from the distribution of insert size of the library. Areas with paired-end evidence of SV are then screened for split-read support. Our analysis only screened for evidence in tandem repeats and deletions, which compose CNVs.

**cn.MOPS**

The cn.MOPS R package provides RD-based CNV calling for WES or WGS data. Briefly, using a cohort of samples, cn.MOPS takes into account the varying number of reads along each chromosome by constructing local read count models. The tool then calls integer copy numbers on segments by using a Bayesian approach to separate technical and biological read count variation from true CNVs.

**CNVkit**

CNVkit is a software for copy number detection, that uses both read depth and nonspecific captured off-target reads to infer copy number evenly across the genome. Since the observed read depth and the true underlying copy number depends on several parameters (e.g. purity, genome ploidy, fraction of the subclonal population), CNVkit initially reports only the estimated log2 copy ratio, but provides several approaches to infer copy number.

**Control-FREEC**

The Control-FREEC tool was developed for detection of allelic imbalances and CNVs. It uses aligned reads to generate a copy number profile, which may then be normalized based on the GC content. For WES data, there is a requirement to use a matched normal sample, and consequently this tool was only applied to the WGS data in this benchmark.

**Manta**

Manta is a structural variant (SV) and indel caller that uses information from mapped paired-end sequencing reads, combining both paired and split-read evidence. Even though Manta can call large-scale SVs, medium-sized indels and large insertions, we have focused here on the last two and ignored SVs, since they are beyond the scope of this paper.

**LUMPY**

LUMPY uses a combined RP/SR/RD approach to call structural variants in WGS data. Using these different kinds of evidence, the LUMPY algorithm probabilistically models structural variant breakpoints. We have focused here on its ability to call CNVs, though the tool also calls other types of structural variants.

**ExomeDepth**

ExomeDepth uses read depth to call CNVs from exome sequencing experiments. The key idea is that the subject exome should be compared to a matched aggregate reference set. As such, this reference set should be made of samples from the same batch and optimized for each exome. In practice, this requirement is rather unpractical, since it is sometimes difficult to build a reference data set for each analysis. As such, we here decided to test ExomDepth on a more unconventional way, building a single (but larger than usual) reference data set. Since ExomeDepth assumes that the CNV of interest is absent from the reference set, there are some caveats that should be kept in mind: i) related individuals should be excluded from the reference; ii) ExomeDepth can miss common CNVs (if they are present in the reference).

**CODEX2**

CODEX2 is an RD-based CNV caller primarily designed for WES and targeted sequencing. It is able to call both somatic and germline CNVs, though we focus here on germline CNVs. Likewise, the tool has different methods for CNV calling with and without negative controls, and we run it in the latter mode.
